# Adipocyte lipolysis abrogates skin fibrosis in a Wnt/DPP4-dependent manner

**DOI:** 10.1101/2021.01.21.427497

**Authors:** A. Jussila, E. Caves, B. Zhang, S. Kirti, M. Steele, V. Lei, E. Hamburg-Shields, J. Lydon, Y. Ying, R Lafyatis, S. Rajagopalan, V. Horsley, R.P. Atit

**Affiliations:** Dept. of Biology, Case Western Reserve University, Cleveland, Ohio, USA; Dept. of Molecular and Cell Biology, Yale University, New Haven, Connecticut, USA; Dept. of Molecular and Cellular Biology, Baylor College of Medicine, Houston, Texas, USA; Department of Medicine, Division of Rheumatology and Clinical Immunology, Pittsburgh, Pennsylvania, USA; Division of Cardiovascular Medicine, University Hospitals, Harrington Heart and Vascular Institute (HHVI), and Case Cardiovascular Research Institute, Department of Internal Medicine and Radiology Case Western Reserve University, Cleveland, OH; Dept. of Genetics and Genome Sciences, Case Western Reserve University, Cleveland, OH; Depts of Dermatology, Case Western Reserve University, Cleveland, OH

## Abstract

Tissue fibrosis in many organs results from altered and excessive extracellular matrix (ECM) protein deposition ^1^. Concomitant with ECM expansion, resident lipid-filled cells including mature adipocytes are lost in human and mouse fibrosis^2-5^, yet the mechanisms that drive mature adipocyte lipid loss and their contribution to tissue fibrosis are unknown. Here, we identify an early, fibro-protective role of mature adipocyte lipolysis driven by Wnt signaling during fibrosis onset. Using chemical and genetic mouse models of skin fibrosis, we show that fibrotic stimuli induce and maintain lipolysis in mature dermal adipocytes. Loss of the lipolytic rate-limiting enzyme adipocyte triglyceride lipase (ATGL)^6,7^ in murine dermal adipocytes exacerbates bleomycin-induced fibrosis development. Adipocyte lipolysis is stimulated in the early stages of Wnt signaling-induced skin fibrosis and by Wnt agonists *in vitro*. Furthermore, deletion or inhibition of the Wnt target gene, CD26/Dipeptidyl peptidase 4 (DPP4) prevented Wnt-induced lipolysis and skin fibrosis in mice. Notably, DPP4 expression correlates with skin fibrosis severity in human patients. Thus, we propose that adipocyte-derived fatty acids and the Wnt-DPP4 axis act as essential regulators of ECM homeostasis within tissues and provide a therapeutic avenue to manipulate fibrosis.

---

Excessive deposition of ECM proteins leads to scarring or fibrosis, inducing tissue stiffening and loss of function in virtually all organ systems, including the skin, adipose tissue, heart, intestine, and lung^1^. Despite its devastating impact on nearly 5% of people worldwide annually^8^, no effective treatment for fibrosis exists. Interestingly, fibrosis occurs concomitantly with a loss of lipid-filled cells in several organs including mature adipocytes in the skin and lipo-fibroblasts in the lung and liver^2-5^.

In the skin, adipocytes compose a distinct layer of dermal white adipose tissue (DWAT) under the skin’s ECM-rich dermal layers, making the skin an excellent system in which to study how lipid-filled cells impact fibrosis development. Accumulation and breakdown of lipids in adipocytes are regulated by the tightly controlled balance between *de novo* lipogenesis, uptake, and breakdown of lipids including lipolysis, lipophagy, and exosomal release of intact lipid^9-11^ (Extended data FigS1.1). We and others have shown that a subset of mature adipocytes undergo dedifferentiation and form myofibroblasts after injury and in bleomycin-induced skin fibrosis in mice^12,13^. In skin repair, this fate transition requires lipolysis and loss of lipid droplets in mature adipocytes in an adipocyte triglyceride lipase (*Atgl)*-dependent manner^14^. However, the role of fatty acids, a product of lipolysis, in fibrosis is unclear. While fatty acids can exacerbate lung fibrosis^15^ and systemic inhibition of lipolysis without fibrotic stimuli can modestly increase homeostatic dermal ECM^16^, fatty acids reduce ECM gene expression in preadipocyte 3T3 L1 cells *in vitro*^17,18^. Together, these studies illustrate the need to better understand the role of lipolysis in initiating and perpetuating fibrosis.

One potential regulator of adipocyte biology during fibrosis is the Wnt signaling pathway. Canonical Wnt signaling through the stabilization of its transducer, β-catenin, drives tissue fibrosis in many organs, including skin^19-21^. Wnt signaling can impact multiple aspects of fibroblast biology including specification, proliferation, migration, myofibroblast formation, and ECM production^22-24^, all of which impact fibrosis pathogenesis. While Wnt signaling can repress adipogenesis^25-28^, the impact of Wnt signaling on mature adipocytes during fibrosis is not known.

## Depletion of lipid from the dermal adipocyte layer is an early event in skin fibrosis

To explore the timing of fibrotic fat loss, skin fibrosis was induced by subcutaneous injection of bleomycin in 6-8 week old mice^29^. While most skin fibrosis models analyze fibrosis induction after 14-21 days, we detected ECM expansion and a 3-fold reduction in Perlipin1^+^ (PLIN^+^) lipid droplet size in mature adipocytes within 5 days of bleomycin treatment (Fig. 1a, b). Lineage tracing of mature adipocytes by tamoxifen-inducible *AdiponectinCreER: mT/mG* reporter^30^ revealed that, following bleomycin injection, GFP+ adipocytes remain in the DWAT region in early stages of fibrosis development, but display smaller or absent PLIN^+^ vesicles (Fig. 1c, Extended data Fig.S1). Examination of dermal adipocytes by electron microscopy^31^ revealed unilocular lipid droplets in control skin whereas bleomycin-injected skin contained adipocytes with multiple smaller lipid droplets (Fig.1d). These data indicate that loss of lipid occurs during skin fibrosis onset, prompting us to confirm whether this was due to lipolysis.

**Fig. 1:**
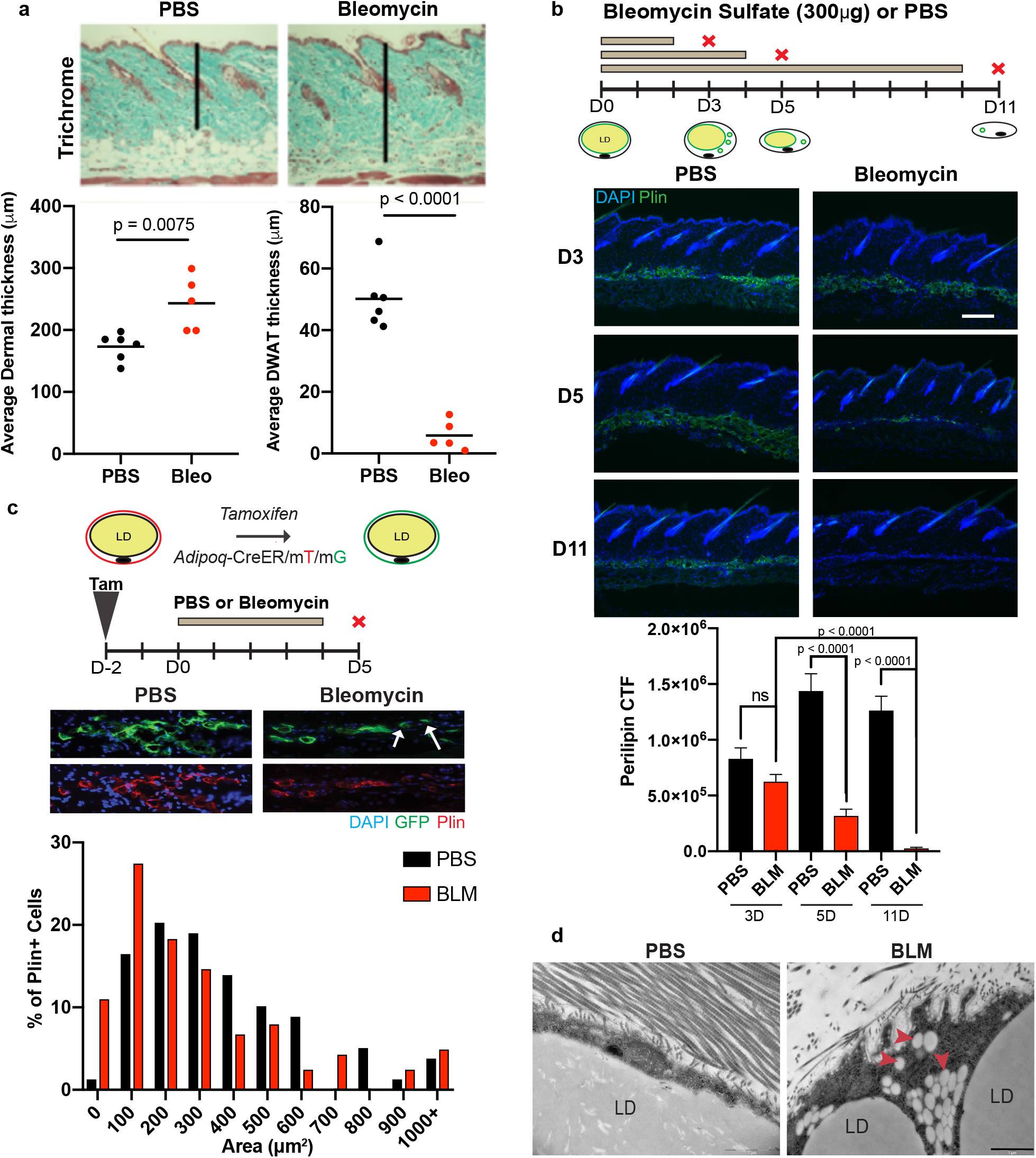
Adipocytes shrink during fibrosis development. **a**,Trichome stained skin sections of WT mice treated with vehicle (PBS) or bleomycin sulfate (BLM) after 5 days. Quantification of dermal and dermal white adipose tissue (DWAT) thickness in these mice. n=5-6 mice for each treatment. **b**, Representative images of perilipin (green) immunofluorescent staining of skin sections from WT mice treated with PBS or BLM for indicated days. Scale bar=200μm. Corrected total fluorescence (CTF) of perilipin immunostaining intensity during time course. n= 2-3 mice per bar. **c**, Genetic lineage tracing of dermal adipocytes in Adiponectin CreER; mTmG mice after 5 days of PBS or BLM treatment. Arrows indicate adipocytes lacking perilipin+ lipid droplets. Histogram of adipocyte size in mice treated with PBS or BLM treatment compared to PBS. **d**, Representative transmission electron microscopy (TEM) images of dermal adipocytes in PBS- and BLM-treated WT mice. LD=adipocyte lipid droplet. Arrowheads indicate lipid vesicles. Scale bar=1μm.

## Inhibition of adipocyte lipolysis exacerbates skin fibrosis development

To test the functional role of adipocyte lipolysis during fibrosis development, we examined mice that specifically lack *Atgl/Pnpla2* in dermal adipocytes^6,7^. Deletion of *Atgl* using *Adiponectin*-*CreER* mice results in severe impairment of FA mobilization during starvation^32^ and after skin injury^14^. We induced skin-specific *Atgl* loss with topical tamoxifen treatment^14^ and subsequently injected mice with bleomycin subcutaneously to trigger skin fibrosis. Bleomycin-injected control (*CreER*^*-*^) mice displayed reduced dermal white adipose tissue (DWAT), however, tamoxifen and bleomycin-injected *Adiponectin*-*CreER; Atgl*^*fl/fl*^ mice retained their DWAT and lipid content (Fig. 2a, b, d). Electron microscopy of dermal adipocytes confirmed unilocular lipid droplets in bleomycin-injected *Adiponectin*-*CreER; Atgl*^*fl/fl*^ (Fig. 2d). Despite retaining dermal adipocyte size and lipid storage, bleomycin-treated *Adiponectin*-*CreER*; *Atgl*^*fl/fl*^ mice displayed precocious dermal thickening (Fig. 2a, b). To explore ECM remodeling in the skin of these mice, we analyzed the levels of unfolded collagen chains using fluorescent collagen hybridizing protein (CHP) in the dorsal skin^33^ (Fig. 2c). We detected increased collagen remodeling throughout the dermis of tamoxifen and bleomycin-treated *Adiponectin*-*CreER; Atgl*^*fl/fl*^ mice compared to bleomycin-injected control mice (Fig. 2c). Together, these data reveal that *Atgl*-dependent adipocyte lipolysis occurs in the early stages of fibrosis and that the early activation of lipolysis during skin fibrosis inhibits ECM expansion during fibrosis induction (Extended data S4.5).

**Fig. 2:**
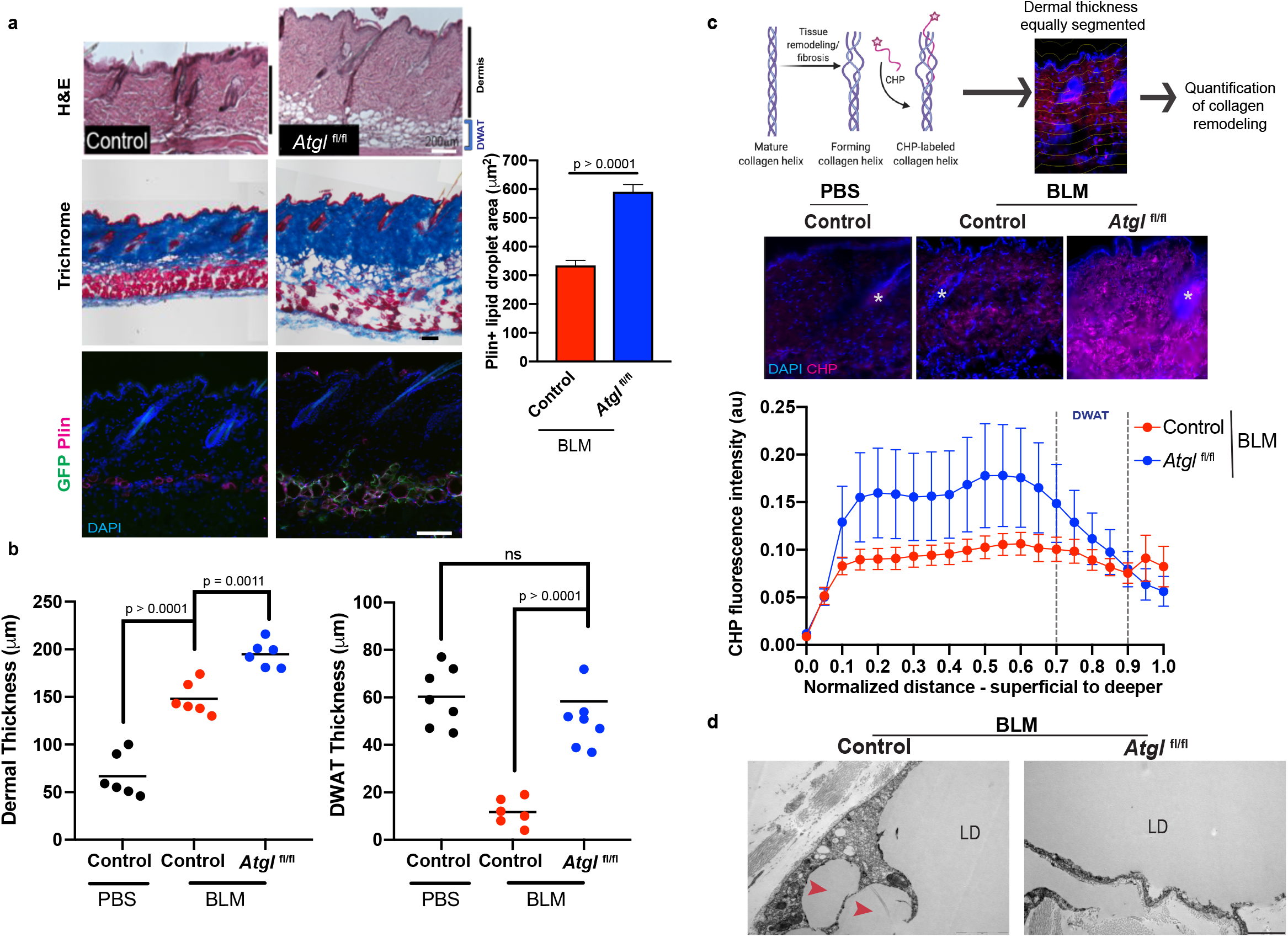
Lipolysis induces adipocyte lipid loss and protects against fibrosis. **a**, Representative images and quantification of skin sections from control (Cre-) and Adiponectin CreER; mTmG; Atglfl/fl (Atglfl/fl) mice after 5 days of control or bleomycin (BLM) treatment stained with indicated dyes (Scale bar=200μm) or perilipin immunostaining (Scale bar=100m). Quantification of Perilipin+ (PLIN+) lipid droplets in control and Atglfl/fl mice. n = 5-7 mice for each genotype. **b**, Quantification of dermal and dermal white adipose tissue (DWAT) thickness in skin sections of control mice treated with PBS or BLM and Atglfl/fl mice treated with BLM. n=5-7 mice per genotype and treatment. **c**, Schematic of pipeline for analysis of collagen remodeling in skin sections. Representative images and quantification of collagen hybridizing peptide (CHP) fluorescent labelling of indicated genotypes and 5 days of control or BLM treatment. n= 3 mice for each treatment. **d**, Representative TEM images of dermal adipocytes from BLM injected control and Atglfl/fl mice. LD=adipocyte lipid droplet. Arrowheads indicate lipid vesicles. Scale bar=2μm.

## The lipolysis axis is stimulated by Wnt signaling in adipocytes

Because Wnt/β-catenin signaling has a key role in fibrosis^19-21,34^ and can impact adipocyte differentiation^25-28^, we analyzed whether Wnt signaling stimulates the lipolytic pathway in skin fibrosis. We induced the expression of stabilized β-catenin, the signal transducer of activated canonical Wnt signaling, in *Engrailed1+(En1)* dermal fibroblasts and adipocyte stem cells^13,22,35^ using *En1Cre/+; R26rtTA/+; TetO-β-catenin/+* (β-cat^istab^) mice (Fig 3a, Extended data Fig. S3.1, S4.3). Dietary doxycycline induced β-cat^istab^ resulted in significant ECM expansion in the dermis and marked DWAT reduction within 10 days (Fig. 3a). β-cat^istab^ throughout the dermis leads to a reduction in DWAT layer thickness (Fig. 3a) and reduced size of individual PLIN^+^ droplets within adipocytes (Fig 3b), despite sustained hair follicle growth (Extended data Fig. S4.3) and associated hair adipocyte enlargement^36^. Interestingly, subsequent withdrawal of doxycycline in β-cat^istab^ for 3 weeks led to rescue of DWAT and dermal thickness (Fig. 3a). Thus, the depletion of lipid within mature dermal adipocytes in mouse skin fibrosis is Wnt signaling-dependent and reversible.

**Fig. 3.**
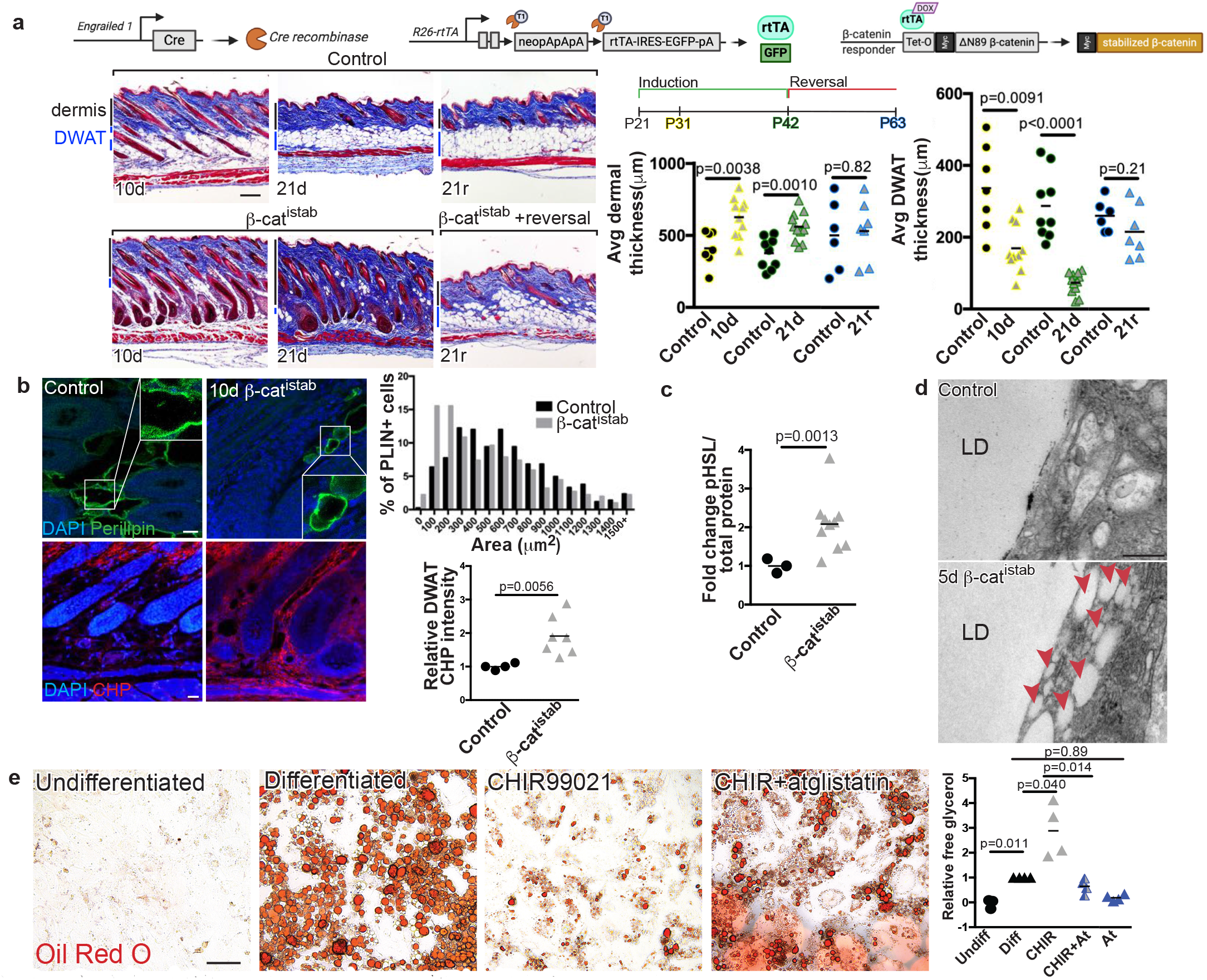
β-cat^istab^ leads to DWAT loss, via dysregulation of lipid metabolism, and collagen remodeling. **a**, Transgenes in doxycycline-inducible/reversible β-catenin(β-cat^istab^) dermal fibrosis model. Fibrosis progression in Masson’s trichrome stained control (top) and β-cat^istab^(bottom) dermis. Black and blue bars indicate dermal and DWAT thickness with quantification of average dorsal dermal and DWAT thickness/mouse. Scale bar=200μm(n=6-11). **b**, Indirect immunofluorescence of PLIN (green) and CHP stain (red) in control and 10d β-cat^istab^ DWAT. Quantification of area of PLIN+ vesicles, 50/mouse (n=9). Relative corrected fluorescence CHP intensity in DWAT. Scale bar=25μm. **c**, Quantification of western blot for pHSL in control and 5d β-cat^istab^ mouse skin relative to total protein (n=3-10). **d**, TEM images of control and 5d β-cat^istab^ DWAT adipocytes. LD=large lipid droplet. Arrowheads=lipid vesicles. Scale bar=200nm. **e**, Primary mouse intradermal adipocyte progenitors untreated(Undifferentiated),treated with adipocyte induction media for 10d and subsequently with maintenance media (Differentiated) Wnt agonist(CHIR99021), or with CHIR99021 and ATGL inhibitor(atglistatin).Scale bar=200μm. Quantification of free glycerol between days 2 and 4 of treatment (n=4).

Next, we examined whether activation of canonical Wnt signaling could induce adipocyte lipolysis by analyzing phosphorylation of Hormone Sensitive Lipase (pHSL) and Perilipin^37^ and glycerol release, an end stage product specific to the lipolysis pathway (Extended data Fig. S1.1). β-cat^istab^ stimulated a nearly two-fold increase in pHSL in β-cat^istab^ skin after 5 days, preceding visible lipid depletion (Fig. 3c). Greater numbers of phospho-Perilipin^+^ adipocytes were also detected (Extended data Fig. S4.3). Electron microscopy confirmed that β-cat^istab^-expressing skin had numerous small intracellular lipid droplets in DWAT adipocytes (Fig. 3d), confirming that lipid dynamics are altered upon Wnt activation. Further, treatment with the Wnt agonist CHIR99021 induced adipocyte lipid loss *in vitro*. Using Oil Red O staining to label lipid content, we observed that treatment of 3T3-L1 differentiated adipocytes and primary mouse dermal adipocytes with CHIR99021 induced loss of lipid in cultured adipocytes (Fig. 3e and Extended data S4.4). Visible lipid depletion was preceded by elevated free glycerol in media, indicating that lipolysis is induced by Wnt activation (Fig. 3e). Glycerol release was abrogated in CHIR99021 treated cells in the presence of an ATGL inhibitor, atglistatin (Fig. 3e), indicating that reduction in lipid content is due to ATGL-dependent lipolysis. Collectively, these data suggest that lipolysis is activated by Wnt signaling and drives lipid loss during the onset of skin fibrosis (Extended data S4.5).

## Wnt-induced DPP4/CD26 is necessary for lipid depletion of dermal adipocytes

To better understand the mechanisms by which Wnt/β-cat signaling promotes fibrosis and adipocyte lipolysis, we analyzed the transcriptome of β-cat^istab^ dermal fibroblasts (GSE 103870)^38^. DPP4 was one of 10 most differentially expressed genes and was highly upregulated (47x, p≤0.005) in β-cat^istab^ dermal fibroblasts^38^. The expression of DPP4 was of particular interest because it is expressed in fibrotic fibroblasts^13^ and its inhibition affects ECM accumulation in injury models^35,39^ and only partially protects from ECM accumulation chemical models of fibrosis^40^, although its role in lipid handling is unknown. DPP4 immunoreactivity was increased in human systemic sclerosis (SSc) and keloids, correlating with SSc disease severity compared to control human skin (Fig. 4a and Extended data Fig. S4.1). Wnt activation induced *Dpp4* mRNA expression in β-cat^istab^ fibroblasts and CHIR99021-treated mouse primary dermal adipocytes *in vitro* (Extended data Fig. S4.2). DPP4 protein is increased in β-cat^istab^ dermis and DWAT *in vivo* and in bleomycin-injected skin (Fig. 4b and Extended data Fig. S4.2). After reversal from β-cat^istab^, *Dpp4* mRNA and DPP4 protein expression levels were restored to control levels *in vitro* and *in vivo*, respectively (Extended data Fig. S4.2). Together, these data demonstrate that DPP4 expression is responsive to Wnt signaling in both dermal fibroblasts and adipocytes.

**Fig. 4.**
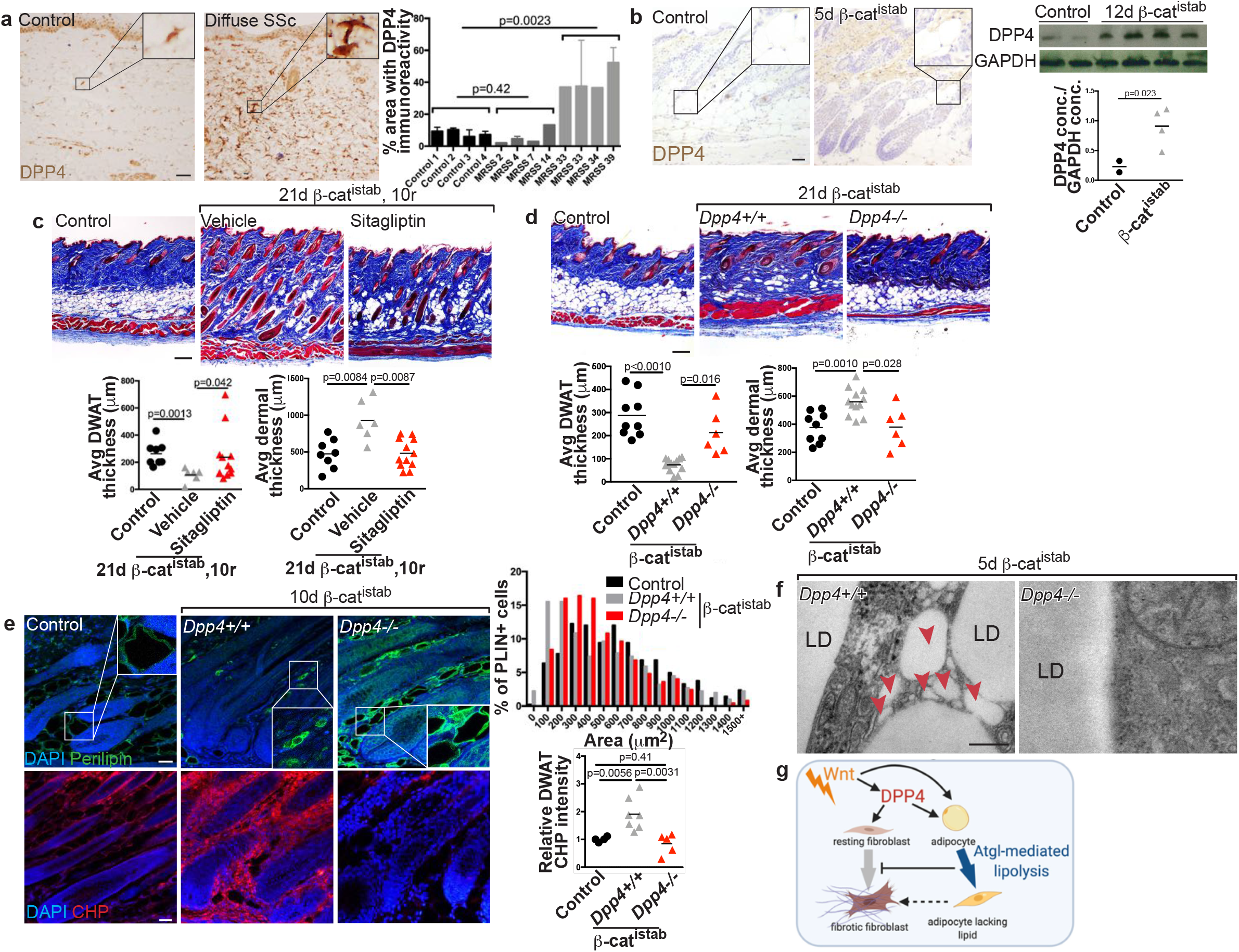
DPP4 mediates lipid loss in β-cat^istab^ dermal fibrosis model. **a**, DPP4 immunohistochemical staining of control and SSc human forearm skin. Scale bar=100μm. Accompanying quantification. **b**, DPP4 immunohistochemical staining on control and 5d β-cat^istab^ mouse skin. DPP4 protein expression relative to GAPDH protein quantity by western blot. Scale bar=100μm. **c**, Masson’s trichrome stained mouse skin from control, 21d β-cat^istab^ and 10d reversal with vehicle or sitagliptin treatment. Scale bar= 200μm. Quantification of dermal and DWAT thickness/mouse (n= 5-12). **d**, Masson’s trichrome stained dorsal skin of control, 21d β-cat^istab^,and *Dpp4-/-* 21d β-cat^istab^. Quantification of dermal and DWAT thickness/ mouse (n= 6-10). Scale bar= 200μm. **e**, Indirect immunofluorescence staining for PLIN (green) and CHP staining (red) in mouse skin from control, 10d β-cat^istab^, and *Dpp4-/-* 10d β-cat^istab^ mice with histogram of PLIN+ vesicle size (n=6-9) and DWAT CHP intensity measurement relative to control. Scale bar= 25μ.m. **f**, TEM images of DWAT adipocytes in *Dpp4+/+* 5d β-cat^istab^ and in *Dpp4-/-* 5d β-cat^istab^. LD=large lipid droplet. Arrowheads= lipid vesicles. Scale bar =200nm. **g**, Workinq model.

Next, we tested if DPP4 inhibition (DPP4i) accelerates recovery from Wnt-induced dermal fibrosis. During early reversal caused by withdrawal of dietary doxycycline for 10 days, β-cat^istab^ skin remained fibrotic and DWAT remained depleted. Treatment of β-cat^istab^ skin during the first 10 days of the reversal phase with a DPP4 inhibitor, sitagliptin, accelerated the recovery of DWAT and dermal thickness (Fig. 4c). *In vitro*, sitagliptin treatment also partially rescued ORO+ lipid droplets in CHIR99021 treated mature adipocytes (Extended data Fig. S4.4). However, sitagliptin co-treatment with CHIR99021 did not fully rescue Wnt-stimulated lipolysis *in vitro*, suggesting that DPP4 function in adipocytes is also likely mediated in part by membrane bound DPP4/CD26, which is not targeted by inhibitors or additional Wnt targets are involved (Fig. 4g).

Subsequently, we tested the hypothesis that Wnt-induced *Dpp4* expression controls adipocyte lipolysis and ECM accumulation *in vivo* by examining whether genetic deletion of *Dpp4* rescues Wnt-induced fibrosis phenotypes. First, we confirmed that Wnt signaling is activated in the dermis and DWAT of *Dpp4*^*-/-*^ ; β-cat^istab^ mice and indeed, we detected nuclear β-catenin in both *Dpp4*^*+/+*^ and *Dpp4*^*-/-*^; β-cat^istab^ mice (Extended Data Fig. S4.3). Strikingly, *Dpp4*^*-/-*^; β-cat^istab^ mice displayed a marked preservation of DWAT, increased PLIN^+^ lipid droplet size, attenuated dermal thickening, in comparison to *Dpp4*^*+/+*^; β-cat^istab^ mice (Fig. 4D and E). Collagen remodeling was significantly diminished in the DWAT and lower dermal regions in *Dpp4*^*-/-*^; β-cat^istab^ (Fig. 4e, Extended Data Fig. S4.3). There was attenuated expression of p-Perilipin and electron microscopy revealed intact unilocular lipid droplets in *Dpp4*^*-/-*^; β-cat^istab^ compared to *Dpp4*^*+/+*^;β-cat^istab^ dermal adipocytes, demonstrating protection from lipolysis (Fig. 4e, 4f, Extended data Fig. S4.3,). Taken together, these data indicate that DPP4 is required for Wnt-induced adipocyte lipolysis and ECM expansion during fibrosis onset and DPP4i can accelerate the recovery from established Wnt-induced fibrosis. These findings resonate with recent work linking DPP4 with obesity, metabolic syndrome, adipocyte dedifferentiation *in vitro*, inhibition of adipocyte differentiation *in vivo*^41-43^, scar formation^13,35^, and fibrosis^39,40,44^.

## Discussion

By combining chemical and genetic models of skin fibrosis development with genetic and pharmacological manipulations (Fig. 4g, Extended data Fig. S4.5), our data unearth molecular mechanisms that drive adipocyte lipolysis and govern ECM homeostasis during fibrosis development. Given the presence of mature adipocytes in the heart and kidney stroma and lipofibroblasts in the lung ^2-5^, the Wnt-Dpp4-adipocyte axis may explain the crucial role of Wnt signaling in concert with other fibrotic stimuli in fibrogenesis of the several other tissues^45^. Furthermore, since fatty acid metabolism^46^, lipid loss^47,48^, and fibroblast activation accompany tumor growth and metastasis^35^, our data may shed light on key mechanisms that impact tissue morphogenetic changes in many disease states.

Our findings also resonate with recent reports that heterogeneous fibroblast populations express DPP4 during skin homeostasis and repair^35,49,50^ and DPPs’ peptidase activity plays a crucial role for inflammation, wound repair, and tumorigenesis^13,51,52^. Together, our data further demonstrate that DPP4 regulates both homeostasis of adipocyte lipid content and fibroblast ECM production to impact tissue fibrosis and promote recovery. We propose that DPP4’s broad substrate repertoire including chemokines and metabolic regulators may drive multiple aspects of fibrosis development and that FDA-approved DPP4 inhibitors may be useful to accelerate clinical treatments for fibrosis prevention and/or recovery.

## Supporting information

Jussila et al., supplemental figures

## Acknowledgements

We would like to contribute all the past and present members of the Atit and Horsley labs that contributed intellectually to this work especially Sarah Ebmeier who initiated this project in the Horsley lab. We thank the CWRU Imaging Core and Yale electron microscopy core. Special thanks to Gregg DiNuoscio, David Buchner, Rodrigo Somoza-Palacios, Jixin Zhong, Meagan Kitt, Emilie Legue and Karl Liem for technical support and advice. The following support contributed to this work: Global Fibrosis Fund (R.P.A), N.I.H-NIAMS R01 AR076938 (V.H. and R.P.A), NIH-NIAMS R01 AR0695505 (V.H.), R01 AR075412, R01 (V.H.), NIH-NIDCR R01 DE18470 (R.P.A), NIH-NICHD R01 HD042311 (J.P.L.), National Institutes of Arthritis Musculoskeletal and Skin Disease grants Scleroderma Center of Research Translation 1P50AR060780 (R.L.), NIH T32 Musculoskeletal Predoctoral Training Grant T32 AR 7505-31 (A.J.), NIH T32 Dermatology Predoctoral Training Grant T32 AR 7569-25 (A.J.), NIH T32 Human Genetics and Genomics Training Grant (5T32HD007149-42) (E.C.), CWRU-SOURCE fellowship (B.Z.), and Arnold and Mabel Beckman Fellowship (SK). Schematics were made on biorender.com.

## Author Contributions

V.H., A.J., E.C. and R.P.A conceived the experimental plan. V.H, R.P.A, A.J., V.L. and E.C. performed the experiments, made the figures, and did the analysis. M.S., E.H-S., S.K., B.Z. and V.L. contributed to data collection. S.K. and B.Z developed and performed machine learning algorithms for quantitative image analysis. J.L, Y.Y., R. L. and S.R. contributed key mouse reagents and human samples.

## Methods

### Mouse handling and lines

*Engrailed1Cre (En1Cre)*^53^; *Rosa26rTA-EGFP*^54^ (Jax Stock 005572) ; *TetO-deltaN89 β-catenin*^55^; *Dpp4*^*-/-*56^; *Adiponectin Cre-ER*^*T2* 57^ (Jax stock 025124); *Atgl* flox^58^ (Jax stock 024278); Rosa26 mTmG reporter^59^ (Jax stock 007676) lines were genotyped as previously described. The contribution of the *En1Cre* lineage cells in the skin has been previously described^13,22,35,60^ . For induction of *TetO-deltaN89 β-catenin-myc* tagged transgene expression in the *En1Cre;R26rtTA* recombined cells, 21-day old (P21) triple transgenic mice were given 6g/kg of dietary doxycycline in rodent chow (Envigo-Harlan) and 2mg/ml doxycycline in water (Sigma) for three weeks. For induction-reversible experiments, P21 mice were first treated with dietary doxycycline until P42 and then switched to regular chow and water. At desired time points, mice were euthanized and dorsal skin was processed for frozen or paraffin sections as previously described^22^. For each experiment, mutants with litter-matched controls were studied. At least two to four litters were used for phenotypic analysis.

Lineage marking of mature adipocytes was achieved using tamoxifen inducible *Adiponectin Cre-ER; Rosa26mTmG* reporter. Topical tamoxifen of 5mg/mL in 100% ethanol is painted directly on the shaved dorsal skin two days prior to bleomycin injection. Conditional deletion of *Atgl/Pnlap2 in mature adipocytes* was done in *Adiponectin Cre-ER; Rosa26mTmG; Atgl*^*flox/flox*^ and verified as previously described^14^ Fibrosis was induced between 6-8 weeks of age in either wild-type or *Adiponectin Cre-ER; Rosa26mTmG; Atgl*^*flox/flox*^ males of C57Bl/6 in bred background. Littermate Cre-negative controls were used for the experiments. Mice are then given daily subcutaneous injection on their upper dorsal region with 300µg of Bleomycin Sulfate (Enzo pharmaceuticals BML-AP302-0010) in 100µL PBS for 3, 5 or 11 days. Bleomycin experiments were done on C57Bl/6 male mice to avoid confounding sexual maturity related differences^61^. Vials of bleomycin sulfate were checked for efficacy prior to experimental treatments. To ensure injections are located to a 0.5in x 0.5 in square, mice are placed under light isoflurane anesthesia prior to injections. 24 hours after the final injection, mice are euthanized and the dorsal skin is collected.

Yale University and Case Western Reserve Institutional Animal Care and Use Committee approved all animal procedures in accordance with AVMA guidelines Protocol 2013-0156, approved 21 November 2014, Animal Welfare Assurance No. A3145-01 at Case Western and Protocol 11248 #A3230-01Yale University.

### Patient Skin samples

Samples were obtained by performing 3 mm punch biopsies from the dorsal mid-forearm of healthy control and systemic sclerosis (SSc) human subjects after informed consent under a protocol approved by the University of Pittsburgh Institutional Review Board. Archived de-identified patient SSc skin samples with Modified Rodnan Skin score (MRSS) were obtained from Scleroderma Center of Research Translation. For human tissue immunohistochemistry, access to archived de-identified keloid tissue was in compliance with the Case Western Reserve University Institutional Review Board for Human Studies.

### Adipocyte cell culture

All data were obtained from primary dermal adipocyte progenitors isolated from wild-type CD1 background P4 dorsal skin. Approximately 1cm^2^ dorsal skin was removed, rinsed in sterile 1x phosphate buffered saline (PBS), minced, and placed in 2mg/mL collagenase (Worthington, LS004196) with 2% bovine serum albumin (BSA) (Fisher, BP1600) and incubated in a 37°C rotating incubator for 45 minutes. Digested skin was filtered through a 70µM cell strainer (cat number) and plated in 60mm tissue culture plastic plates. Cells were passaged 2-3 times when they achieved approximately 80% confluence. Adipocyte differentiation was stimulated with adipocyte induction media (AIM) (DMEM (Thermofisher, 11995065) containing glucose, pyruvate, 10% fetal bovine serum (FBS), 100µM indomethacin, 1µM dexamethasone, 500µM 3-isobutyl-1-methylxanathine (IBMX), and 10µM insulin) for 8-12 days. Duplicate cultures were kept in media containing only glucose and pyruvate. Differentiated adipocytes were enriched by trypsinizing (0.25% Trypsin EDTA (Thermofisher: 25200056)) and re-plating on a 12-well plate for treatment. Some differentiated adipocytes were kept in maintenance media only (DMEM with glucose, pyruvate, FBS, and insulin), or with additives such as 7µM CHIR (Cayman, 13122), 40µM atglistatin (Sigma Aldrich, SML1075-5MG), and/or 20µM sitagliptin (Cayman, 13252). All media was prepared fresh and changed every 2^nd^ day. Following 4 days of treatment, media was collected from each well and free glycerol was quantified using Free Glycerol Reagent (Sigma-Aldrich F6428-40ML). Data was plotted relative to untreated control mature adipocytes from the same mouse and analyzed using a paired *t*-test with Welch’s correction. Following 8 days of treatment cells were rinsed and fixed for 1 hour in 10% neutral buffered formalin. They were then stained with Oil Red O (ORO) (Sigma Aldrich, O0625-25G) for 10 minutes and rinsed with water before they were imaged on a Leica DMi8 inverted microscope. Camera positions within wells were held constant between plates and each well was photographed in 2 non-overlapping regions. Images were then quantified using Cell Profiler™ (See below) to estimate coverage and average coverage per condition was displayed relative to untreated control cells and analyzed with a paired *t*-test with Welch’s correction. Experiments were repeated on 3-4 biological replicates.

### Histological staining and morphometrics

Dorsal mouse skin from mice of various ages (p26, p32, p42, p68) was isolated, fixed in 4% PFA, and equilibrated in 25% sucrose for cryosectioning. Dorsal back skin was isolated from 6-8-week-old mice. It was directly frozen in OCT for cryosectioning at 14

μm. Alternatively, dorsal skin pieces were drop-fixed 10% neutral buffered formalin for 1 hour at 4 degrees and then processed for paraffin sectioning at 7μm. Sections were stained with Masson’s trichrome for mature Collagen I expression or hematoxylin and eosin according to standard protocols. Brightfield images were captured with an Olympus BX60 microscope and Cell Sens entry software and Zeiss AX10 scope and Zen 2.6 pro software. Dermal thickness, DWAT thickness, and adipocyte number were quantified with measurement tool in Fiji/Image J software. Data represent the average thickness in three different regions in 5-10 non-overlapping fields/mouse^12,19^.

### Immunohistochemistry and Immunofluorescence

For Figures 1 and 2 and Extended Data S1.1, frozen tissue was sectioned at a thickness of 14µM for immunofluorescence staining. Slides were fixed for 10 minutes in 4% paraformaldehyde (PFA) prior to staining. For Figures 3 and 4 and Extended data Fig 3.1, 4.1-4.5, drop-fixed paraffin tissue was embedded and sectioned at 7µM. Paraffin sections were deparaffinized and washed with 1xPBS. They underwent antigen retrieval in citrate buffer (10 mM Tri-Sodium Citrate dyhydrate, 0.05% Tween-20, pH 6.0) for 15 minutes at 93°C in a water bath. Following 10% normal goat serum block, with 0.05% Tween-20 or 0.3% Triton for 1 hr at room-temperature, tissue was incubated with appropriate primary antibody overnight at 4°C. Primary antibodies for GFP (Abcam ab13970, 1:1000), β-Catenin (BD Biosciences, 1:250), myc-tag (Cell Signaling 7D10, 1:50), perilipin (Abcam ab3526, 1:500; Abcam ab61682, 1:500), p-perlipin1 (Vala Sciences, 4856), and CD26/DPP4 (Abcam ab28340, R&D: AF954) were used for brightfield immunohistochemistry or immunofluorescence as previously described^19,55,62^. After three washes in PBST (PBS+ 0.3% Triton or 0.05% Tween), species-appropriate secondary antibodies conjugated to biotin (Vector) or Alexa-fluor (Thermo-Fisher) were used. Nuclei were counterstained with hematoxylin or DAPI (1:2,000) before mounting in Fluoroshield (Sigma). Negative controls were used to confirm antibody specificity.

*Imaging:* Brightfield Images were takes at room temperature. Brightfield images in Figure 1 and 2 were obtained using Zeiss AX 10 upright scope equipped with a digital camera (Hamamatsu, ORCA-Flash4.0) and a 10x Objective (Zeiss Plan-APOCHROMAT). Brightfield images of Masson’s Trichrome staining in Figure 3 and 4 and extended data E3.1, 3.2, 3.3 were taken using a Olympus BX60 microscope with a digital camera (DP70, Olympus) with Cell Sens Entry software (Ver. 1.5, © Olympus Corporation 2011) with a 4x objective (Olympus UPIanFI 4x/0.13) . Exposure was held constant between controls and experimental group. Brightfield images of Oil Red O stained cells in Fig. 3E and Extended data figure S4.4 were imaged on an inverted widefield Leica Dmi8 microscope with a digital camera and a 10x objective (HC PL FLUOTAR 40x/0.60, Dry, FWD=3.3-1.9mm).

Immunofluorescence images in Figure 1 and 2 and extended 1.1 were obtained on the Zeiss AX 10 upright scope equipped with a digital camera (Hamamatsu, ORCA-Flash4.0) and a 20x Objective (Zeiss Plan-APOCHROMAT). Immunofluorescence images in Figure 3B and 4E were imaged on an inverted confocal Leica TCS SP8 gated STED 3x microscope (DMI6000, Leica), using a 40x oil immersion objective (HC PL APO 40x/1.30 NA CS2, Oil, FWD=0.24 mm) detection by PMT detector and/or hybrid detectors and Leica LAS X software. CHP images were taken at 40x magnification on an inverted widefield Leica Dmi8 microscope. Max projections were generated and assembled using Fiji/ImageJ and analyzed in Cell Profiler. Images were processed and merged using Adobe Photoshop and laid out Adobe InDesign or Illustrator.

### Image treatment and analysis

#### Adipocyte perilipin+ cross-sectional area

For Figure 1 and 2, 20x images on the Zeiss AX10 scope were analyzed. For Figure 3 and 4, 40X confocal images were analyzed in FIJI ImageJ. With FIJI’s polygon selection tool, at least 50 adipocytes were counted per mouse from non-overlapping fields. The areas of these identified adipocytes were binned in a histogram generated in GraphPad Prism.

#### Corrected total Perilipin fluorescence

20x images of WT mice stained with Perilipin were analyzed using FIJI. The DWAT layer, which contained perilipin was outlined. The area and perilipin fluorescence intensity were measured. Three full-thickness fields of the dermis (perilipin negative) were measured for fluorescent intensity to determine the average mean background fluorescence. Corrected total fluorescence was measured as perilipin intensity - (area * average background).

#### ORO Pipeline

Prior to feeding a batch of 4x images through the Cell Profiler™ pipeline, an undifferentiated control ORO image was first used for white-balancing using Adobe Photoshop’s image processor. The configurations of this white-balancing action were input into a white-balancing script and the full batch of images were white balanced with the script. Next, the images were loaded into Cell Profiler™ and placed through an *Image Math* module (Operation: Invert, Multiply the first image by: 1.5) to invert the image. The inverted images were converted into grayscale images using a *Color To Gray* module (Conversion method: Split) and split into RGB channels. Each of the 3 channels were run through a series of *Correct Illumination Calculate* and *Correct Illumination Apply* illumination correction modules. The grayscale “Green” and “Blue” channels were added together in another *Image Math* module (Operation: Add) to more clearly show the ORO-stained objects on a black background. An *Identify Primary Objects* module (Threshold smoothing scale: 1.3488, Threshold correction factor: 1.0, Lower and upper bounds on threshold: 0.0, 1.0, Size of adaptive window: 50) configured with an adaptive, 2-class Otsu thresholding method was used to detect ORO-stained objects based on intensity. Finally, a *Measure Image Area Occupied* module was used to output both the area covered by ORO-stained objects as well as the total area of the image in squared pixels.

#### DPP4 Pipeline (SSc)

Prior to feeding a batch of 4x images through the Cell Profiler™ pipeline, a human forearm-skin DPP4 control image was first white-balanced using Photoshop’s image processor. The configurations of this white-balancing action were input into a white-balancing script to be used on the entire batch. These white-balanced images were loaded into Cell Profiler™. A *Crop* module was used to manually select a rectangular, representative ROI in each image. Color deconvolution was completed using an *Unmix Colors* module to separate the images into Hematoxylin and DAB channels. A *Reduce Noise* module (Size: 7, Distance: 11, Cut-off distance: 0.045) was used on the DAB channel. Unwanted objects including blood vessels and hair follicles were manually identified through an *Identify Objects Manually* module. These identified objects were transformed into a binary image via a *Convert Objects to Image* module. The area of these unwanted objects was determined and recorded by a *Measure Image Area Occupied* module. Using a *Mask Image* module (invert the mask: yes), a mask of the unwanted objects was applied into the original ROI rectangle. The resulting ROI, with unwanted objects masked out, was thresholded by a *Threshold* module via an adaptive, 3-class Otsu thresholding method in which the intermediate intensity objects were classified as foreground objects (Threshold smoothing scale:0.0, Threshold correction factor: 1.15, Lower and upper bounds on threshold: 0.15 & 1.0, Size of adaptive window: 50). The area of these threshold identified DPP4+ objects as well as the total ROI area recorded by a *Measure Image Area Occupied* module. These recorded area data were output by an *Export To Spreadsheet* module. The relevant total ROI area was determined by subtracting the unwanted area from the total ROI area. The area covered by DPP4+ objects was divided by the relevant total ROI area to give the percent coverage of DPP4+ per image.

<u>B-CHP pipeline: adapted from https://onlinelibrary.wiley.com/doi/pdf/10.1111/exd.13457</u> 40X B-CHP images were loaded into Cell Profiler. Each original image was first split into grayscale versions of its RGB channels using a *Color To Gray* module (Conversion method: Split). An *Identify Objects Manually* module was used to trace and select an ROI composed of the entire skin under the epidermis and above the panniculus carnosus. A *Convert Objects To Image* module was used to convert the ROI into a binary image. A *Closing* module (Structuring element shape, size: disk, 50) was applied to this binary ROI to fill in any gaps left by the tracing performed in *Identify Objects Manually*. Using the shape of this corrected binary ROI object, a sequence of *Morph* (performed operation: distance) and *Image Math* modules are used to generate an intensity-based distance map of the ROI based on distance from the epidermis. *Rescale Intensity* (Rescaling method: Divide each image by the same value) modules then established how far down subsequent layering modules would extend. Divisor values for *Rescale Intensity* modules were changed for each image to the maximum thickness (pixel length) from the epidermis to the panniculus carnosus. Using the epidermis distance map, *Threshold* modules (Threshold strategy: global, thresholding method: manual, Threshold smoothing scale: 1, Manual Threshold: increased from 0 to 1 in 0.05 increments), established binary regions of increasing distance from the epidermis. *Image Math* modules (Operation: subtract) were then used to subtract each thresholded binary region from the preceding region to establish preliminary layers. This generated 20 preliminary layers spanning the ROI. *Erode Image* modules (Structuring element shape, size: square, 10) were used to slightly shrink each layer and ensure no overlap. An *Image Math* module (Operation: add) was used to add eroded layers to produce a full map of the layers. A sequence of identification and conversion modules was used to produce a binary image containing the area occupied by the layers. Using an *Identify Objects Manually* layer, hair follicles and all area including and under the *panniculus carnosus* were selected from the original B-CHP image. Using a sequence of *Mask Image* and *Mask Objects* modules, these unwanted objects were masked out of the binary image containing the area occupied by the layers. This new masked binary image was composed of the area defined by the layers minus the area of the unwanted objects. The new binary image was converted back into objects using a *Convert Image To Objects* module. These objects were masked over the original layers objects to generate the final map of the layers excluding unwanted areas. The red intensity from the original red channel was then calculated per final layer using a *Measure Object Intensity* module. A *Measure Object Size Shape* module was used to calculate the area of each final layer. The final layers were then overlaid on the original image to ensure correct functioning of the pipeline. The results were output using an *Export To Spreadsheet* module.

### Transmission Electron Microscopy analysis

Mice were anesthetized with ketamine and xylazine prior to perfusion. Mice were fixed by transcardial perfusion with 4% PFA according to standard protocols^31,63^. Samples were then submitted to the Yale EM core for processing and imaged as previously described^31^. Briefly, hardened blocks were cut using an ultramicrotome (UltraCut UC7; Leica). Ultrathin 60-nm sections were collected and stained using 2% uranyl acetate and lead citrate for transmission microscopy. Carbon-coated grids were viewed on a transmission electron microscope (Tecnai BioTWIN; FEI) at 80 kV. Images were taken using a CCD camera (Morada; Olympus) and iTEM (Olympus) software.

### RNA extraction, and qRT-PCR analysis

Total RNA was extracted from cultured cells using Trizol reagent (Thermo Fisher: 15596026) and processed for qRT-PCR analysis with 4ng of cDNA as previously described {Hamburg-Shields:2015kd}. *Axin2* and *Dpp4* mRNA quantities were measured relative to *Hprt* using Taqman master mix (Thermofisher, 4304437) and probes (Thermofisher, Mm00443610_m1, Mm00494549_m1, Mm03024075_m1). Relative mRNA quantities were determined using an Applied Quantstudios Biosystems 3 PCR System. All samples were normalized to *Hprt* gene expression, and results are expressed as the fold change of Ct values relative to controls, using the 2−ΔΔCt formula. Complete qRT-PCR data was depicted in univariate scatter plots as recently described ^64^. Statistical significance was determined by two-tailed, unpaired Student *t*-test with Welch’s correction in GraphPad Prism software.

### Western Blot analysis

Western blot was performed on flash frozen dorsal mouse skin samples. Samples were mechanically dismembrated, suspended in 1xRIPA buffer (Cell Signaling 9806S) and sonicated. 10-30µg protein was loaded per well. When stain free gel was used (pHSL blot), it was activated prior to transfer to PVDF membrane and total protein was detected before primary antibody incubation. Otherwise transfer to PVDF was performed without activation or total protein detection (DPP4 blot). Blots were incubated in 5% milk and 3% BSA block respectively for 1 hour at room temperature followed by incubation with α-DPP4 (R&D: AF954) or α-phospho-HSL (Cell signaling: 4139) primary antibody overnight at 4°C followed by species appropriate secondary antibody. Protein from stain-free blots (pHSL) were visualized on a Bio-Rad Chemidoc™ gel imager and data was graphed relative to total lane protein calculated in Image Lab software. Otherwise (DPP4), blot was developed on film, stripped, and re-probed with GAPDH loading control. Densitometry was calculated in Fiji and graphed relative to GAPDH loading control.

### Statistical analysis

Sample size was determined based on published studies and no statistical method was employed. The experiments were not randomized. Due to the nature of the genetic manipulations, the authors were not blinded to allocation of animals for the experiments. The authors were blinded during results of the analyses. Individual data points on graphs represent the average value per mouse of 2-12 biological replicates (depending on measurement). Normality in the spread of data for each experiment was tested using Shapiro-Wilk test in GraphPad Prism software. Significance values for data sets

displaying normal distributions were calculated by unpaired Student *t*-test (two-tailed, unequal variances) with Welch’s correction in Prism software. Additionally, one-way analysis of variance (ANOVA) was performed on Prism to compare dermal and DWAT thickness between PBS control, BLM-Cre^-^ control and BLM *Atgl*^*fl/fl*^. Paired t-tests are performed where appropriate (*in vitro* only). Significance for non-normal distributed data were calculated using the Mann-Whitney *U*-Test in Prism software. For graphs with individual data points, each point represents the average of one mouse. Error bars represent standard error. All p values are included on the graphs and p values less than

0.05 are considered statistically significant.

## Data availability

The data sets analyzed during the current study are available in the GEO repository (GSE 103870).

