## Supplementary material for "Adipocyte lipolysis abrogates skin fibrosis in a Wnt/DPP4-dependent manner": Jussila et al., supplemental figures

**a**

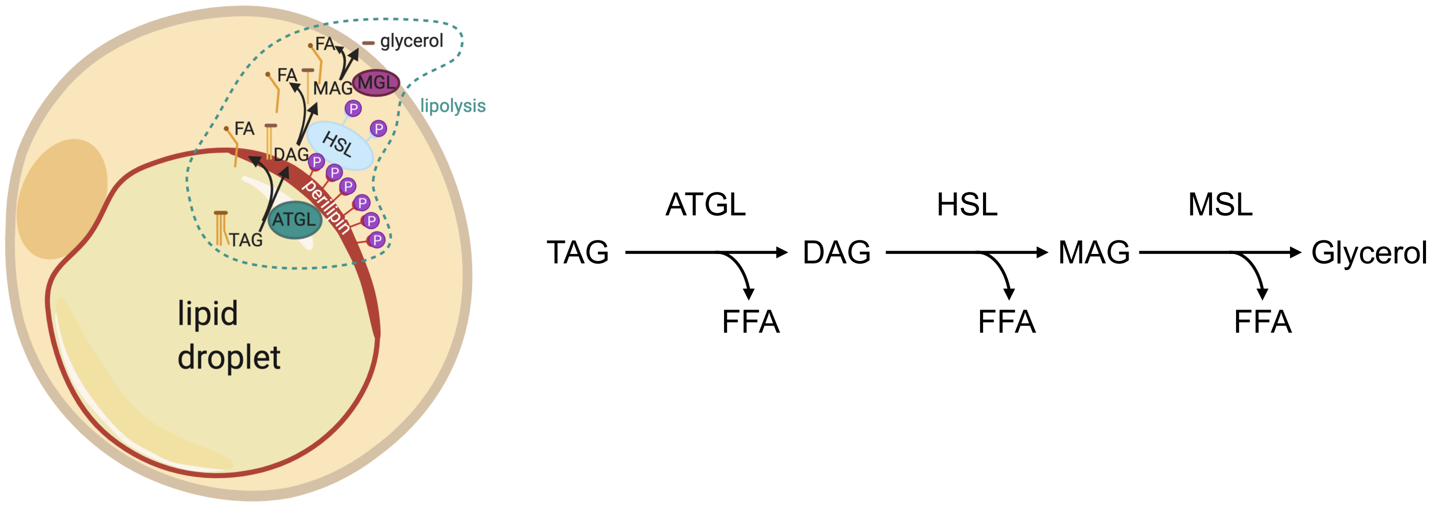


**b**

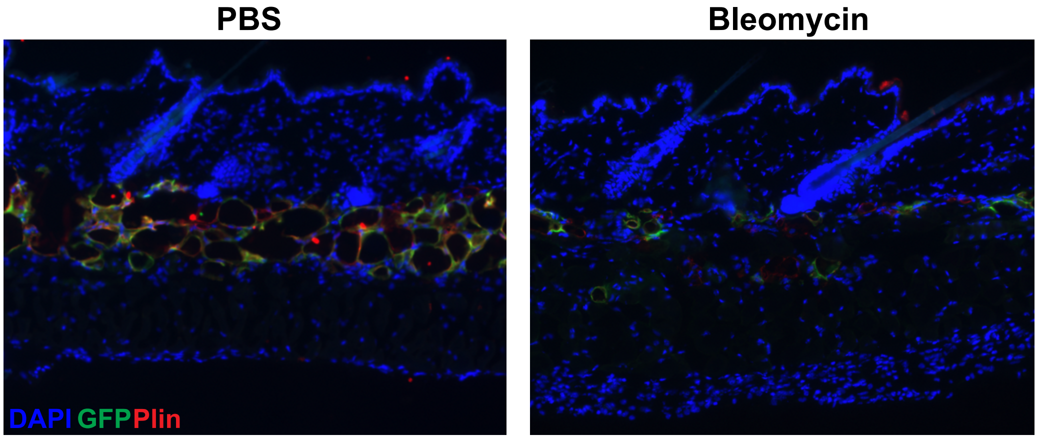


**Fig. S1.1: At 5 days of Bleomycin treatment, membrane GFP positive adipocytes do not appear in the dermis**
**a**, Schematic of the ATGL-dependent lipolysis pathways, breaking down triglycerides (TAG) to three free fatty acids (FFA) and a glycerol.
**b**, mTmG mice were injected for 5 days with PBS or BLM. GFP marks dermal adipocytes. GFP+ cells are only present in the DWAT layer after 5 day bleomycin treatment.

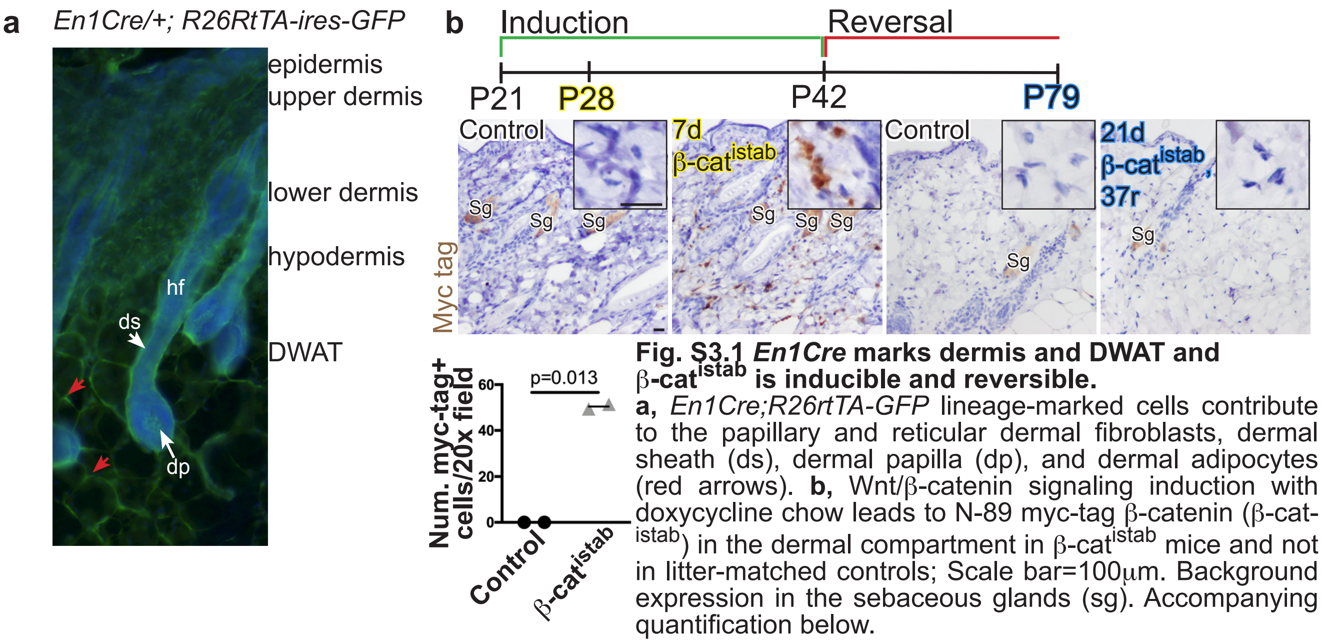


**Fig. S3.1 *En1Cre* marks dermis and DWAT and β-cat$^{istab}$ is inducible and reversible.**
**a,** *En1Cre;R26rtTA-GFP* lineage-marked cells contribute to the papillary and reticular dermal fibroblasts, dermal sheath (ds), dermal papilla (dp), and dermal adipocytes (red arrows). **b,** Wnt/β-catenin signaling induction with doxycycline chow leads to N-89 myc-tag β-catenin (β-cat$^{istab}$) in the dermal compartment in β-cat$^{istab}$ mice and not in litter-matched controls; Scale bar=100μm. Background expression in the sebaceous glands (sg). Accompanying quantification below.

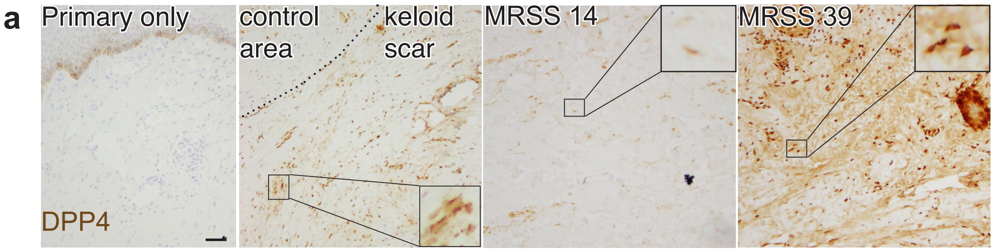


**Fig. S4.1 DPP4 expression is elevated in keloid involved region and in high MRSS scored human skin.**

**a,** IHC with $\alpha$-DPP4 antibody from left to right: Primary only control, human keloid biopsy (dotted line indicates boundary of keloid determined by histology), low MRSS (14) scored human forearm biopsy sample, high MRSS scored (39) human forearm biopsy sample. Scale bar= 100$\mu$m.

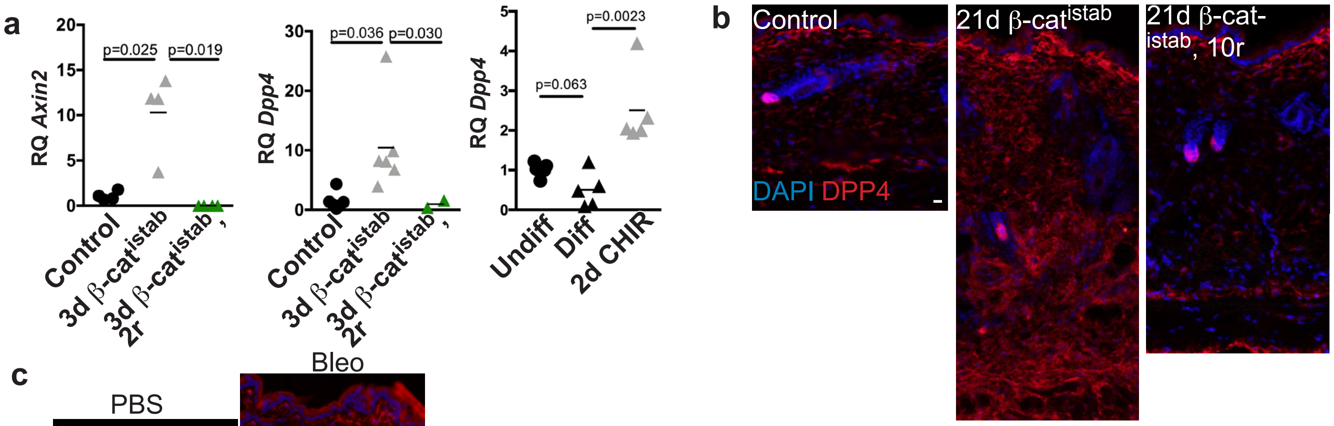


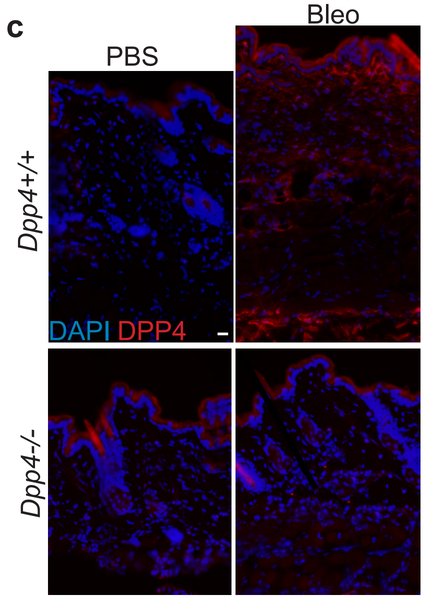


**Fig. S4.2 Dpp4 is Wnt responsive *in vitro* and *in vivo* and increased in a chemical fibrosis model.**
**a,** *Axin2* (left) and *Dpp4* (center) mRNA quantities relative to *Hprt* mRNA quantity in dermal primary cells following β-cat$^{istab}$ and reversal. MRNA quantity of *Dpp4* relative to *Hprt* in cultured dermal primary cell-derived fibroblasts and adipocytes with or without 48 hours Wnt agonist treatment (7μM CHIR99021) (right). **b,** *En1Cre-* control (left), 21d β-cat$^{istab}$ (center), and 21d β-cat$^{istab}$, 10d reversal (right) dorsal dermis subjected to IF with α-DPP4 antibody; Scale bar=25μm **c,** Dorsal dermis of *Dpp4+/+* (top) and *Dpp4-/-* (bottom) mice injected subcutaneously with PBS (left) and bleomycin (right) subjected to IF with α-DPP4 antibody; Scale bar=25μm.

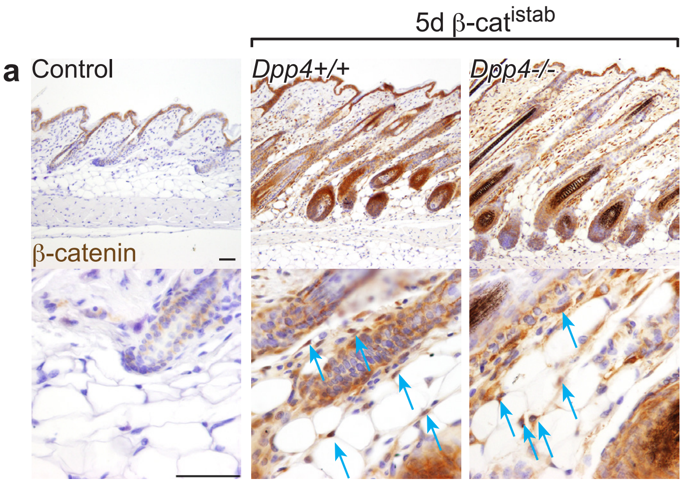


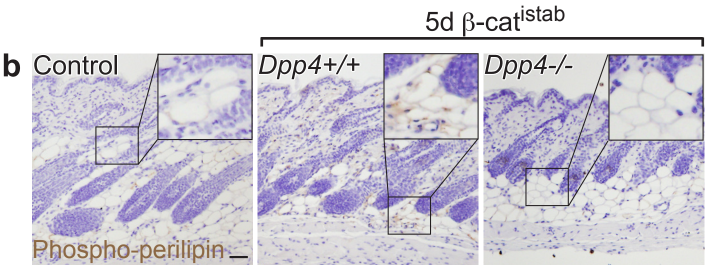


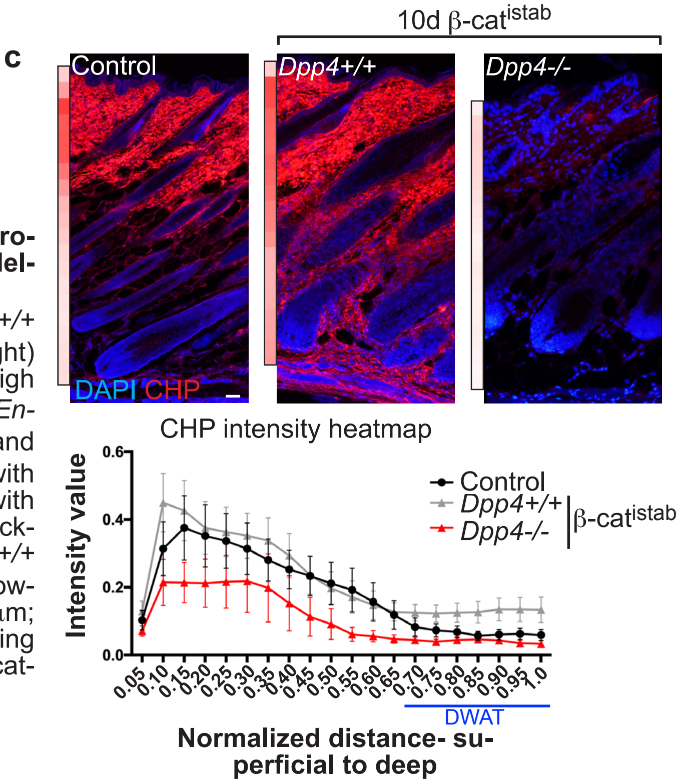


**Fig. S4.3 *Dpp4-/-* mice achieve β-cat$^{istab}$, but are protected from fibrotic lipolysis and DWAT CHP remodeling.**

**a,** β-catenin IHC in representative control (left), *Dpp4+/+* 5d β-cat$^{istab}$ (center), and *Dpp4-/-* 5d β-cat$^{istab}$ (right) dorsal dermis. Blue arrows indicate adipocytes with high β-catenin signal. Scale bar=100 μm. **b,** Age-matched *En-1Cre-* control (left), *Dpp4+/+* 5d β-cat$^{istab}$ (center), and *Dpp4-/-* 5d β-cat$^{istab}$(bottom) mouse dorsal dermis with IHC stain for p-plin protein. Photos are taken at 10x with high powered field inset. Scale bar=100μm. **c,** Full thickness dorsal dermis of En1Cre- control (left), 10d *Dpp4+/+* β-cat$^{istab}$ (center), and 10d *Dpp4-/-* β-cat$^{istab}$ (right) following CHP stain with heatmap bars on left. Scale bar=25μm; CHP intensity map below, values thresholded by staining batch, with SEM bars; Approximate DWAT location indicated with blue bar.

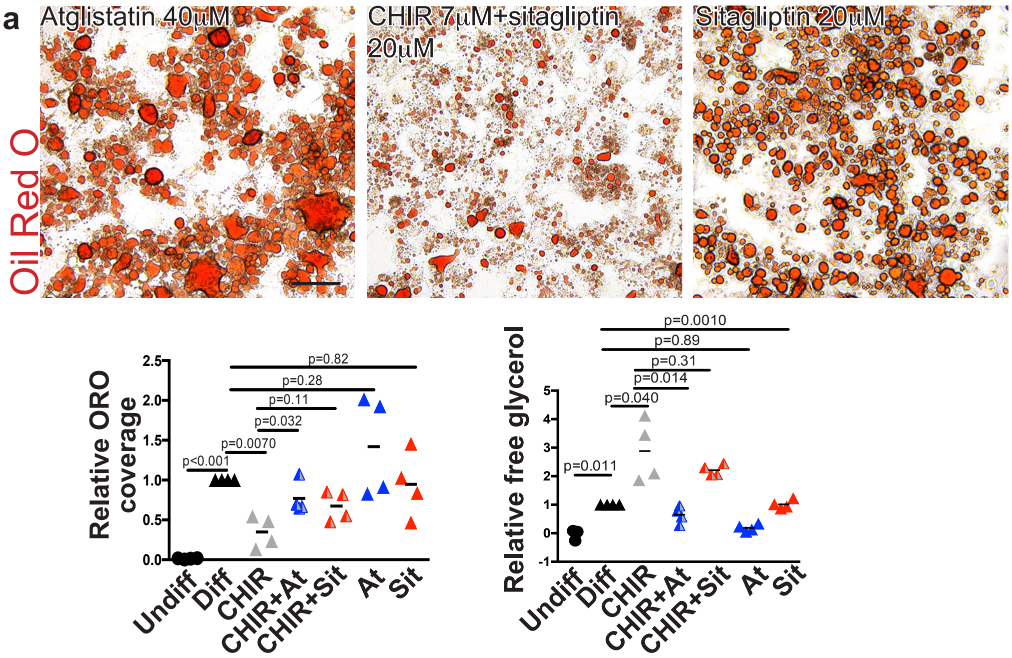


**Fig. S4.4 Primary dermal adipocytes are protected from lipid loss, but not from lipolysis by DPP4$^{I}$.**

**a,** Primary intradermal adipocyte progenitor cells treated with differentiation media followed by 40μM atglistatin (left) 7μM Wnt agonist, CHIR99021, and 20μM DPP4 enzymatic inhibitor, sitagliptin, (center), or treated with differentiation media, followed by 20μM sitagliptin (right); Scalebar= 200μm; Average Oil Red O area covered quantification (2 fields per well, 4 biological replicates, and 2-4 technical replicates per sample per treatment) (left bottom). Average free glycerol release between 2d and 4d of previously described treatments (right bottom).

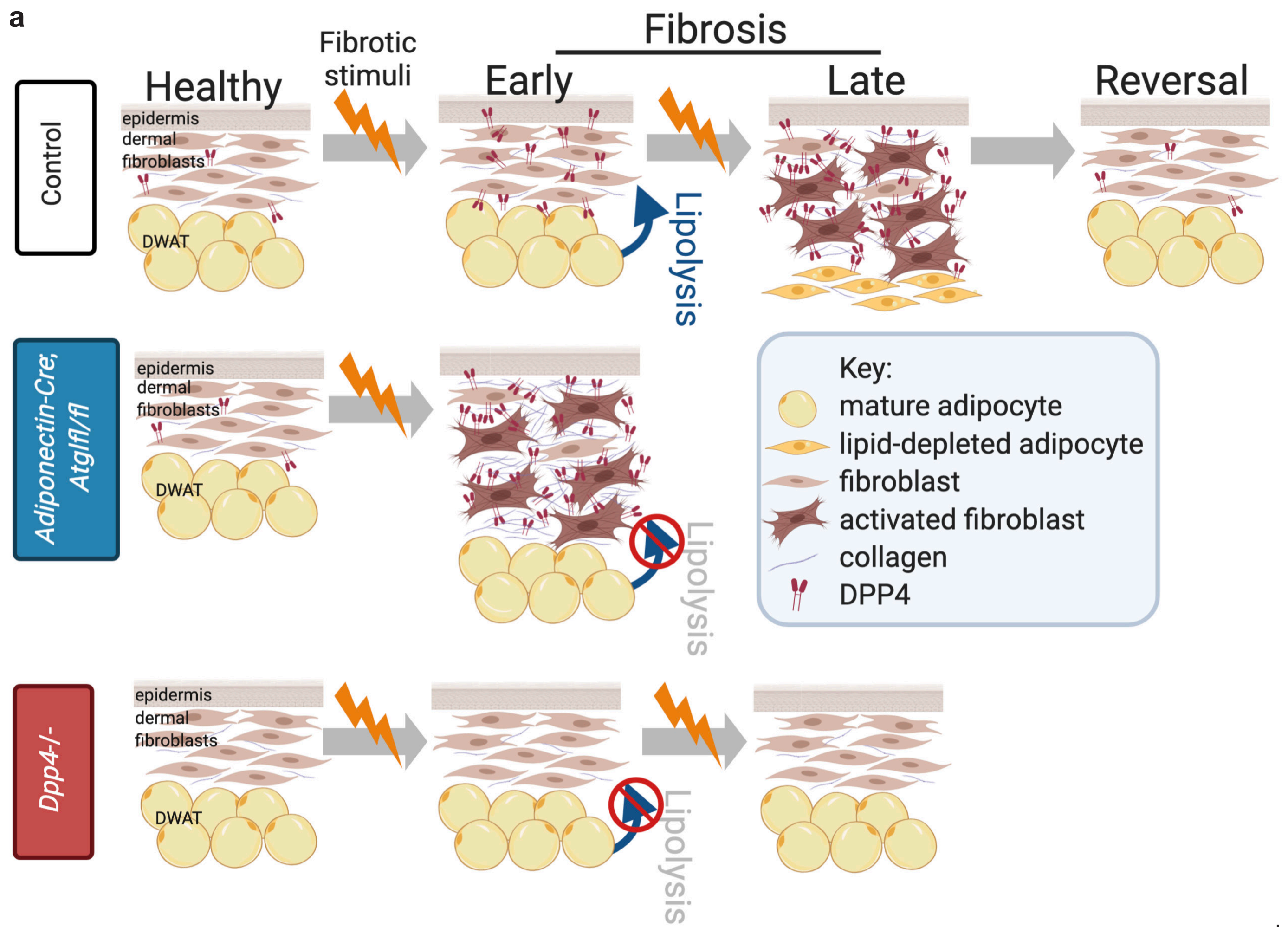


**Fig. S4.5 Summary schematic. a,** ATGL-mediated lipolysis occurs early in bleomycin and $\beta$-cat$^{istab}$ models. Dermal *Atgl* mutants have preservation of DWAT, but greater dermal thickening and collagen remodeling. DPP4 is Wnt-responsive and increased in mouse models and *Dpp4-/-* mice have preservation of DWAT and reduced dermal thickening and collagen remodeling despite $\beta$-cat$^{istab.}$
